# Production of HPV16 capsid proteins in suspension cultures of human epithelial cells

**DOI:** 10.1101/2020.10.28.359273

**Authors:** Dirce Sakauchi, Erica Akemi Kavati Sasaki, Fernanda de Oliveira Bou Anni, Aurora Marques Cianciarullo

## Abstract

Human papillomavirus (HPV) infection is a leading cause of morbidity and mortality in women worldwide. The virus is associated with benign warts and a broad spectrum of malignancies, including cervical cancer, considered a disease of high clinical relevance, especially in developing countries. In this study we developed the production of recombinant proteins HPV16 L1 and HPV16 L2 in human cells in suspension (293-F), which were transiently co-transfected with the pUF3/L1h and pUF3/L2h vectors. Expressions of recombinant HPV16 L1 and L2 capsid proteins was detected by laser scanning confocal microscopy and flow cytometry. Both proteins were identified intracellularly in the nucleus and cytoplasm of cells. The presence of these heterologous proteins and VLPs formation were detected by transmission electron microscopy (TEM) through colloidal gold immunolabeling and negative staining. Cell extracts containing recombinant proteins were purified by affinity chromatography and immunization of Balb/c mice with the formulation HPV16 L1/L2 VLPs containing adjuvant was able to induce higher titer of anti-HPV16 L1, when compared to HPV16 L2 antibodies by indirect ELISA assay. These data indicate that transient expression in 293-F cells was efficiently established. The results are promising for obtain recombinant proteins of the HPV capsid for future studies involving human papillomavirus, as well as to contribute for the development of other vaccine strategies for prevention against HPV.

## Introduction

Cervical cancer is the most relevant consequence of human papillomavirus (HPV) infection, being considered the most common cancer among women worldwide, representing a major global challenge for public health.

The HPV involvement in cervical cancer cases is well established, as well as has been associated with lower prevalence with other cancers: anogenital, head, neck, oropharynx and cutaneous cancers [1, 2]. Nowadays there are over 200 genotypes of human papillomavirus already identified [3] causing about 5.2% of all human cancers [4]. At least 19 HPV types are considered of high oncogenic risk [5] with more than 70% of cases of cervical cancer worldwide associated with HPV16 and HPV18 [6, 7]. The HPV16 is considered the most prevalent serotype, detected in more than 50% of cases [8].

Human Papillomavirus is a non-enveloped virus containing an icosahedral capsid formed by two structural proteins: the major L1 and minor L2. The L1 coat protein can self-assemble into virus-like particles (VLPs), present in current HPV vaccines, which are expressed in *Saccharomyces cerevisiae* (quadrivalent and nonavalent Gardasil - Merck) or insect cells (bivalent Cervarix - GlaxoSmithKline) [3, 9]. The HPV quadrivalent and bivalent vaccines induce high immunogenicity and effectively protect against its specific serotypes present in the vaccines, with limited cross-protection against other HPV [10, 11]. Other possibility of production would be the use of the viral capsid protein L2, which can also produce VLPs when combined with L1. The L2 is the smallest protein in the viral capsid and contains highly conserved sequences for broader protection against various types of HPVs.

Previous studies shown several strategies for recombinant protein expression and production of HPV VLP, using bacteria, yeast, insect and mammalian cells [12, 13]. This study related the production of structural proteins HPV16 L1/L2 using human epithelial cells (293-F) suspended in medium without fetal bovine serum (SFB). Human epithelial cells (HEK293) have been one of the most widely used in recombinant protein production because these cells are easy to grow and have been considered of high transfection efficiency for protein production [14, 15]. In this study, the 293-F cells derived from lineage HEK 293 adapted to grown in suspension were used for the production of recombinant proteins of HPV. In addition to ethical issues, the medium in the absence of serum has several advantages in terms of eliminating potential risk of contaminant minimizing animal components, meeting regulatory requirements, and simplifying the process of protein purification.

The use of suspension cell culture system without animal-derivate components is an alternative strategy for the production of recombinant proteins, contributing for diverse applications. In this study we describe viral capsid protein expression, characterization and evaluation of immune response in a murine model.

## Materials and methods

### Expression of HPV16 L1/L2 recombinant proteins for VLPs production

Human cells adapted in suspension (FreeStyle™293-F, Invitrogen) were cultured in serum-free medium (Freestyle™293 Expression Medium, Invitrogen, USA) at 37°C with 5% of CO_2_ in an orbital shaker at 120 rpm. In the present study, the pUF3/L1h and pUF3/L2h gene expression vectors used were constructed under the control of the human cytomegalovirus promoter (pCMV), in which are expressed the respective humanized HPV16 L1 and L2 genes, and also contain the ampicillin resistance gene. Both vectors were provided by Prof. Dr. Martin Müller from DKFZ-Heidelberg, Germany. For vector amplification, *Escherichia coli* DH5α was grown at 37°C in Luria Broth (LB) medium (1% tryptone, 0.5% yeast extract, 1% sodium chloride) supplemented with 100 μg/ml ampicillin. The plasmid DNA purification was performed with AxyPreplasmid Miniprep assay kit (Axygen Biosciences, USA) according to the manufacturer instructions and plasmid concentration was determined on a Nanodrop 100 spectrophotometer (Thermo Scientific, USA).

For expression of the recombinant proteins, HEK 293-F cells in suspension at a density of 2×10^6^ cells/mL were transiently transfected with pUF3/L1h or co-transfected with pUF3/L1h or pUF3/L2h vectors, using 293 fectin reagent (Invitrogen, USA) according to the manufacturer instructions. Following the transfection, the cells were incubated at 37°C with 5% CO_2_, under agitation at 120 rpm. A maturation time is required for capsids production and stabilization [16]. Briefly, cell lysate was incubated overnight at 37°C, and then chilled, adjusted to 0.8 M NaCl, clarified by centrifugation at 2,000 x *g* for 15 min at 4°C. Expressions of the HPV16 L1/L2 heterologous proteins and the formation of VLPs were analyzed by immunofluorescence, ultrastructural immunocytochemistry and Western-blot.

### Cell lysis and purification of HPV16 L1/L2 proteins

Suspension of co-transfected 293-F cells was washed with phosphate- buffered saline (PBS) and centrifuged at 200 x *g* for 10 min. Cells pellet was resuspended in lysis buffer containing protease inhibitor (Sigma Aldrich™, USA) and lysed by vortex agitation followed by incubation on ice. After centrifugation at 500 x *g* for 10 min., the lysate clarified was precipitated with ammonium sulfate to 45% saturation, under slow agitation at 4°C during 30 min. The solution was then centrifuged at 12,000 x *g* at 4°C for 10 min and the precipitate was resuspended in PBS, dialyzed against PBS/Tween-80 at 4°C for 72 h. The dialysate was applied in HiTrap™ Heparin HP chromatography (GE Healthcare, Sweden) column, equilibrated in PBS/0.01% Tween-80 plus 0.3M NaCl (3 column volume) and the L1/L2 proteins were eluted with 1M NaCl in PBS/0.01% Tween-80. Fractions (1-5) were collected and analyzed by SDS-PAGE and Western-blot.

### Protein assay

Protein concentration was determined by the bicinchoninic acid protein assay kit (BCA-Pierce, USA) using bovine serum albumin (BSA, Sigma-Aldrich) as standard.

### SDS-PAGE and Western-blot

Samples of proteins were separated by 10% SDS-PAGE (Sodium Dodecyl Sulphate PolyAcrylamide Gel Electrophoresis), under reducing conditions and stained with silver nitrate or transferred to nitrocellulose membrane (Trans-blot transfer membrane BioRad, USA). For Western-blot, membranes were blocked overnight with 5% non-fat milk in PBS containing 0.05% Tween 20 (PBS-T), washed in PBS and incubated with anti-HPV16 L1 (BD Pharmingen, USA), anti-HPV16 L2 (Santa Cruz Biotechnology, USA) or anti-L1/L2 VLP IgG for 1h at room temperature. After washing in PBS-T, the membranes were incubated with goat anti-mouse IgG peroxidase-conjugated (Sigma-Aldrich) diluted in 5% non-fat milk in PBST for 1 h. Then, the membranes were washed in PBS and the L1 and L2 proteins were detected using Novex ECL HRP Chemiluminescent Substrate Reagent Kit (Invitrogen).

### Indirect immunofluorescence

To detect HPV16 L1/L2 intracellular protein expression, transfected cells were washed three times in PBS and fixed with 2% paraformaldehyde (PFA) at 4°C for 1 h. The cells were washed again in PBS and blocked in PBS containing 5% bovine serum albumin (BSA) for 1 h. Following three washes, the cells were incubated with primary antibodies anti-HPV16 L1 (BD Pharmigen) and anti- HPV16 L2 (provided by Prof. Dr. Richard B. Roden, from John Hopkins University, USA), diluted in PBS-T containing 0.5% BSA for 2 h at room temperature. Then, the cells were washed again in PBS and incubated with the respective secondary antibody Alexa Fluor 488 goat anti-mouse IgG (Molecular Probes) for L1 protein detection, or Alexa Fluor 633 goat anti-rabbit IgG for protein L2, diluted in PBST containing 1.5% BSA for 1 h at room temperature. After three washes, the cells were mounted on Mowiol (Calbiochem, USA), covered with coverslips and analyzed in an LSM 510 Meta laser scanning confocal microscope using LSM 5 Image software (Carl Zeiss, Germany).

### Flow cytometry

For quantitative flow cytometric analysis, the suspension of cells with different times of co-transfection (6, 12, 24, 36, and 48h) was washed in PBS and centrifuged at 800 rpm for 5 min. Next, cells were fixed in 2% PFA for 1 h at 4°C, washed again and incubated with 3% BSA in PBS/0.05% Tween 20 for 1 h. After washing in PBS, cells were incubated with the same primary antibodies anti-HPV16 L1 or anti-HPV16 L2 and secondary antibodies used in immunofluorescence assay. Then, cells were washed again and analyzed in BD FACS Canto™ II cytometer using BD FACS Diva software (BD Biosciences, USA). For transfection efficiency analysis, fluorescence intensities were measured for 10,000 events for each experiment.

### Ultrastructural immunocytochemistry

Co-transfected cells in suspension were washed in PBS and in 0.1M phosphate buffer. Cells were then fixed in 3.5% sucrose, 4% paraformaldehyde (Sigma-Aldrich, USA) and 1% glutaraldehyde in 0.1M phosphate buffer. After washes, cells were dehydrated in a gradual ethanol series (50%, 70% and 80%), pre-embedded in 2:1 acrylic resin (LR White, Electron Microscopy Sciences, USA) and 70% ethanol, embedded in 100% LR White acrylic resin for 1 h, then in 100% resin overnight. The resin was changed again and samples were distributed in gelatine capsules and polymerized at 60°C for 48 h.

Ultrathin sections were mounted on nickel grids pre-coated with 2% parlodion and carbon [18]. For immunogold labeling, the thin sections were incubated first with primary antibodies anti-L1 and anti-L2. Then they were incubated with secondary antibodies goat anti-mouse IgG complexed with gold colloidal particles (10-nm) for L1 and goat anti-rabbit IgG conjugated with gold particles (15-nm) for L2, diluted in PBS containing 0.01% Tween 20 and 1.5% BSA for 1 h at room temperature. The grids were rinsed with PBS and water, contrasted with 2% uranyl acetate and examined in Zeiss EM109 transmission electron microscope operated at 80 kV, with digital camera system for images capture Megaview G2, Olympus Soft Imaging Solutions.

### Negative staining

Purified VLPs samples were adsorbed to carbon-coated grids and negatively stained with 2% uranyl acetate in aqueous solution. Grids were allowed to air-dry prior to examination with a Zeiss EM 109 transmission electron microscope operated at 80 kV, with digital camera system for images capture Megaview G2, Olympus Soft Imaging Solutions. Micrographs were taken with various magnifications [12, 13, 18].

### Immunization of animals

Balb/c mice of six to eight weeks-old in groups of six animals were subcutaneously immunized three times at two weeks intervals with 20 μg HPV16 L1/L2 VLPs alone (protein) and adsorbed (formulation) with 200 μg aluminum hydroxide as adjuvant. The control group was inoculated with saline. Serum samples were collected two weeks after the last immunization and analyzed by ELISA and Western-blot assays.

### Enzyme-linked immunosorbent assays (ELISA)

The presence of anti-HPV16 L1 and HPV16 L2 IgG in sera of immunized mice were determined by indirect ELISA assays. Briefly, the HPV16 L1 and HPV16 L2 proteins were expressed in 293-F cells as previously described. Ninety-six well plates were coated with 100 μL of HPV16 L1 or HPV L2 proteins purified by affinity chromatography and size-exclusion chromatography, respectively (Sephacryl^TM^S300) and diluted in carbonate buffer (pH 9.6) at 4°C. The plates were washed again with PBS-T and blocked with 3% BSA in PBS for 2h at 37°C. The plates were washed three times and incubated with dilutions of the mice sera (pool) diluted in PBS-T with 1% BSA for 2 h at 37°C. After washing step, 100 μL of the secondary antibody (goat anti-mouse IgG peroxidase-conjugated from Sigma) was added for 1 h at 37°C. The reaction was developed with 100 μL of the orthophenylenediamine (OPD) in citrate phosphate buffer (pH 5, 0) and hydrogen peroxide for 15 min at room temperature. Optical density was measured at 492 nm on a Multiskan Ex plate reader (Labsystems Uniscience). Statistical analysis was performed using the Prism software (GraphPad) and the statistical difference between groups was analyzed by Student's *t*-test where p<0.05 was considered statistically significant.

## Results

### Expression and localization of recombinant proteins

The expression of recombinant proteins HPV16 L1 and HPV16 L2 was evaluated in 293-F cells transiently transfected with pUF3/L1h and pUF3/L2h vectors.

In order to identify the highest expression of recombinant protein, kinetic experiments were performed using confocal laser microscopy and flow cytometry. The kinetics of expression of the HPV16 L1 and HPV16 L2 proteins revealed that cells showed positive reaction at 6 hours after transfection, using anti-HPV16 L1 and L2 antibodies by immunofluorescence assays (Fig. 1A). The maximal expression was observed around 48 hours post-transfection by confocal laser scanning microscopy (Fig.1E). In these experiments it was possible to observe the expression of intracellular proteins with double immune labeling of HPV16 L1 in green and HPV16 L2 in red, distributed in the nucleus and cytoplasm of the cells (Fig 1A - 1E). Both proteins were not detected in the control cell (Fig. 1F).

**Fig. 1.**
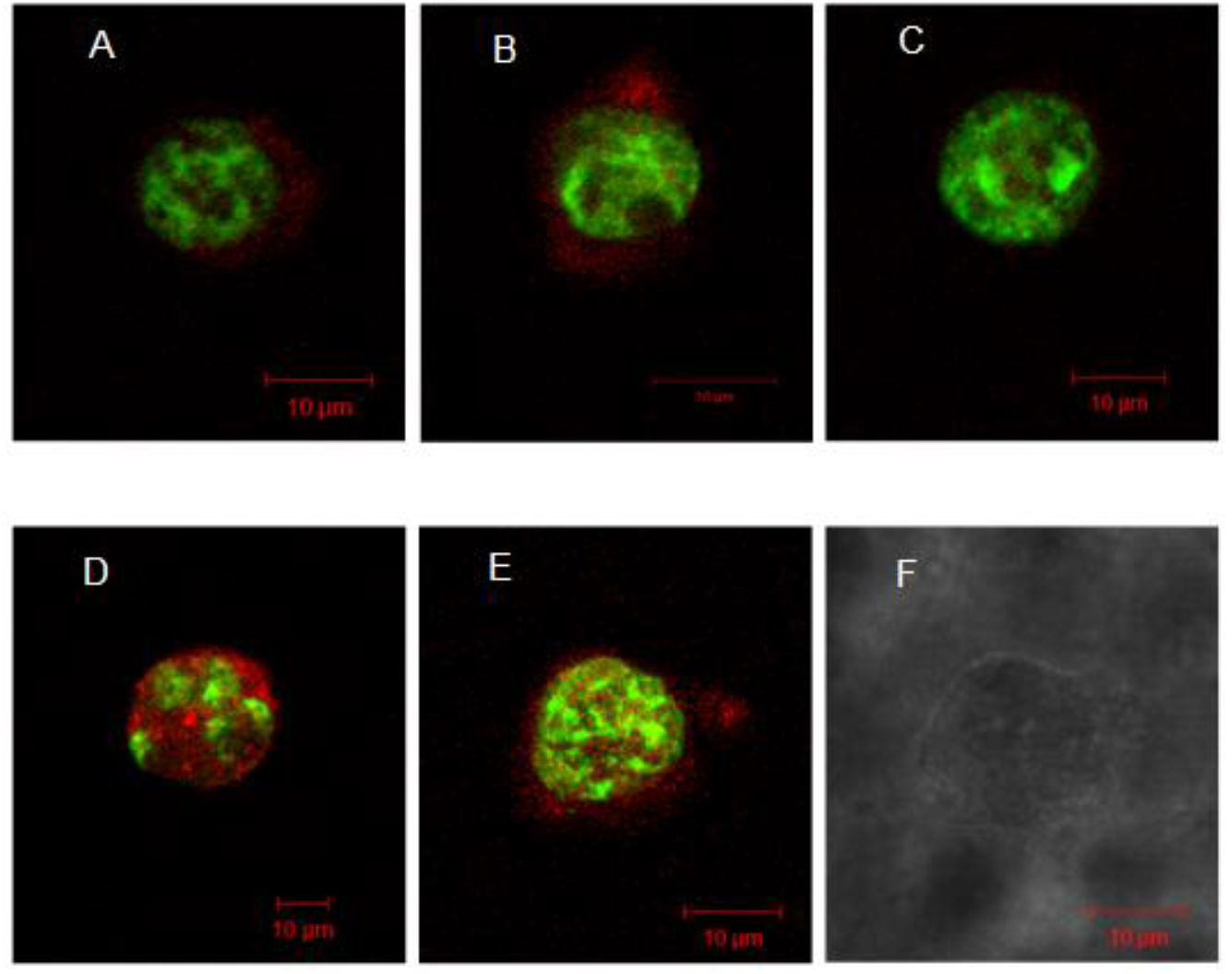
Analysis of the recombinant HPV16 L1 and L2 proteins expression. 293-F cells were co-transfected with plasmids pUF3/L1h and pUF3/L2h, obtained at different times (a = 6h, b = 12h, c = 24h, d = 36h, e = 48h, f = control) post-transfection and visualized by confocal laser scanning microscopy. The protein L1 (green) was detected with anti-HPV16 L1 and secondary Alexa Fluor^®^ 488 anti-mouse IgG produced in goat. HPV16 L2 protein (red) revealed with anti-HPV16 L2 antibody and Alexa Fluor^®^ 633 goat anti-rabbit IgG (L2). Magnification: Objective C-Apochromatic 63xs / 1.4 Oil. Bar= 10μm

High expression of these recombinant proteins was also detected by flow cytometry, showing that approximately 83% of cells were expressing the L1 and L2 proteins after 48 hours of cellular transfection.

Electron microscopy of co-transfected cells showed localizations for both the L1 and L2 proteins in the same cellular regions, inside cytoplasm and nucleus of cells (Fig. 2). Both recombinant proteins were identified using anti-HPV16 L1 and anti-L2 antibodies, followed by secondary antibodies complexed with 10-nm and 15-nm colloidal gold particles, respectively.

**Fig. 2.**
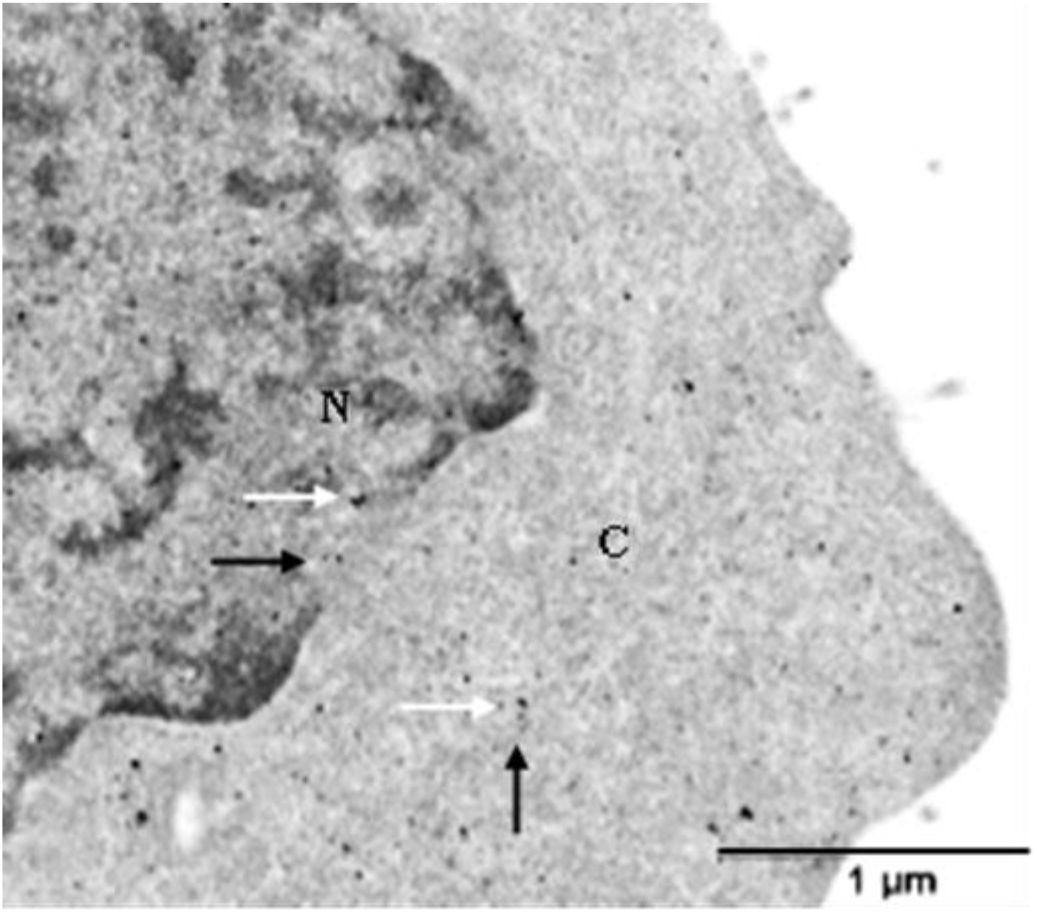
Detection of L1 and L2 proteins by transmission electron microscopy (TEM). Ultrathin section of co-transfected cell processed by special procedure and immune labeling with anti-L1 and anti-L2 antibodies conjugated to 10 nm (L1) colloidal gold particles (black arrow) and 15 nm (L2) (white arrow) dispersed in the nucleus (N) and cellular cytoplasm (C). Magnification= 17,000x. Bar=1μm

Ultrastructural immunocytochemistry assays showed evidence of recombinant proteins self-assembling similar to virus-like particles (VLPs) with heterogeneous sizes varying approximately between 20 to 55 nm in diameter. Furthermore, it was also possible to observe the presence of HPV VLP immune labeling for proteins L1 (10 nm gold) and L2 (15 nm gold) (Fig. 3A). Structured HPV16 L1/L2 VLPs and purified were also examined by transmission electron microscopy, showing an icosahedral shape composed by L1 pentamers, at about 50 nm in diameter (Fig. 3B).

**Fig. 3.**
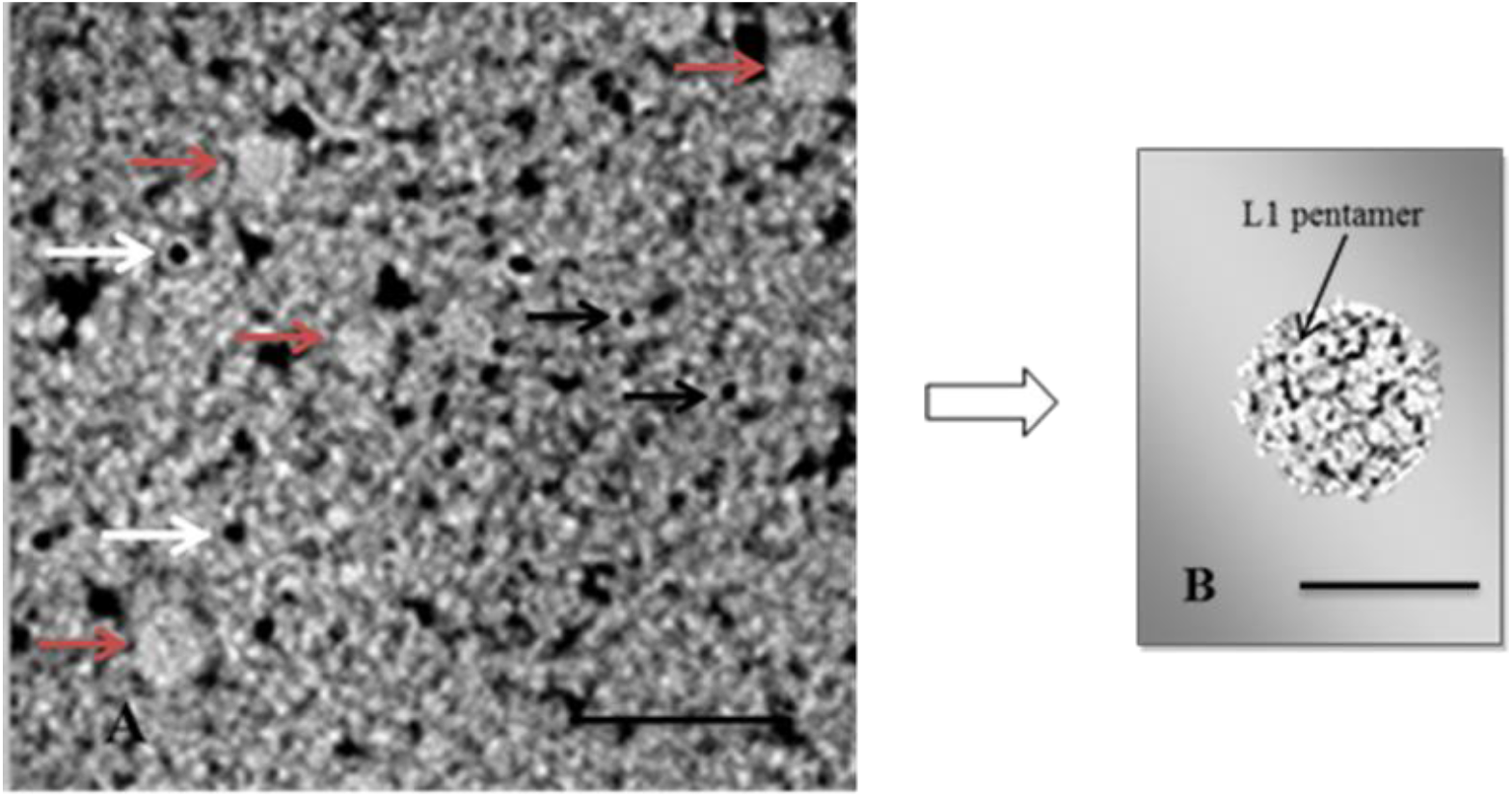
Expression of recombinant proteins HPV16 L1 and L2 by the ultrastructural immunocytochemistry and negative staining analysis. (**A)**Co-transfected cell immune labeling with anti-L1 and anti-L2 antibodies, conjugated to colloidal gold particles of 10-nm (L1) (black arrows), and 15-nm (L2) (white arrows). VLPs formation by self-assembling of L1 and L2 proteins (red arrows). Original magnification = 30,000x. Bar = 100 nm. (**B**) Transmission electron micrograph of HPV16 L1/L2 VLP by negative staining is showing a structured and stabilized HPV16 capsid, composed by L1 and L2 proteins. Examined in Zeiss EM109 Transmission Electron Microscope operating at 80 kV with digital camera Megaview G2, Olympus Soft Imaging Solutions. Original magnification = 50,000x. Bar = 50 nm

### Characterization of proteins L1/L2

SDS-PAGE analysis of purified recombinant L1/L2 proteins (fractions 1-5) obtained from affinity chromatography demonstrated the presence of protein L1 with expected molecular mass of approximately 55 kDa and other band with around 72 kDa, similar to protein L2. Lower molecular mass proteins were also visualized, probably resulting from proteolytic degradation (Fig. 4A).

**Fig. 4.**
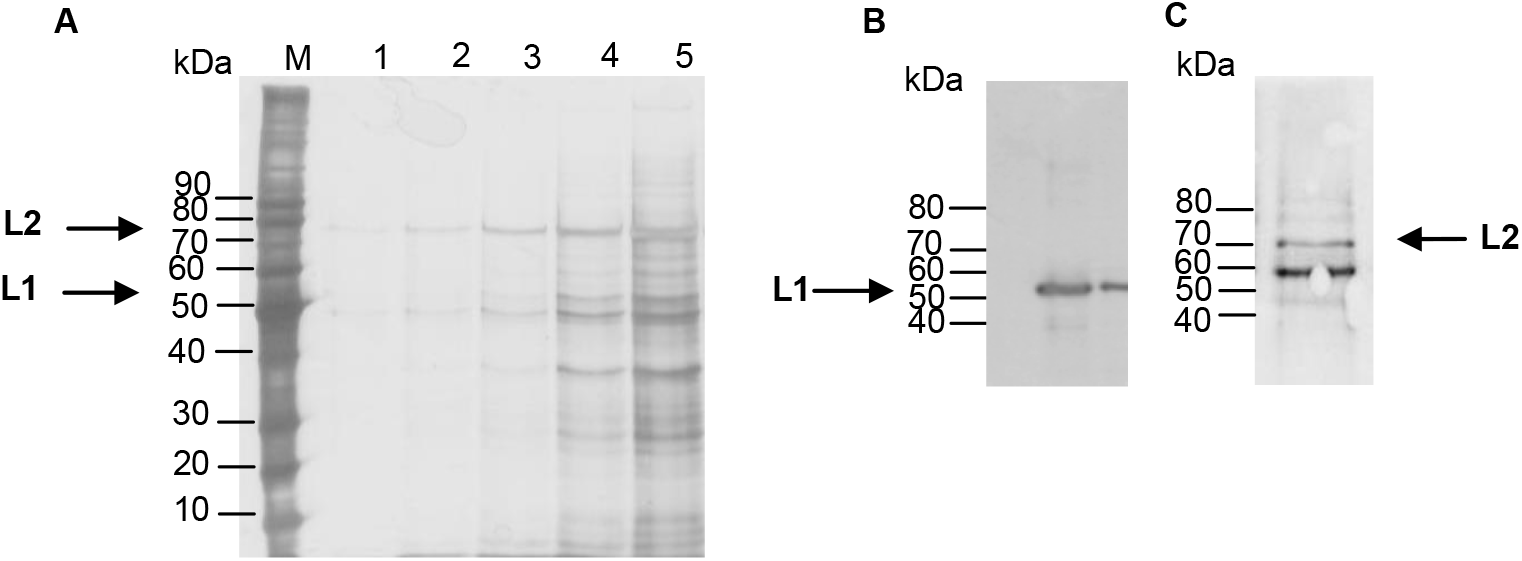
Characterization of HPV16 L1/L2 proteins. (**A**) 10% SDS-PAGE. Molecular mass standards (M); Fractions (a1-a5) eluted from affinity chromatograph; (**B**) Western-blot of purified proteins (pool 1-5). L1 protein (~ 55kDa) revealed by anti-HPV16 L1 (**C**); L2 protein (~72kDa), detected by monoclonal anti-HPV16 L2 antibody, followed by secondary antibody goat anti-mouse IgG peroxidase

Both intracellular proteins were identified by Western-blot, showing a band similar to the L1 protein in purified L1/L2 samples (pool 1-5) that was specifically recognized by the monoclonal antibody anti-HPV16 L1. In addition, the band corresponding to protein L2 was immune reactive for the anti-HPV16 L2 monoclonal antibody (Santa Cruz Biotechnology, USA) (Fig. 4B).

### Analysis of anti-HPV16 L1/L2 VLPs antibodies

Balb/c mice were immunized subcutaneously with HPV16 L1/L2 VLPs (protein) or adsorbed (formulation) with aluminum hydroxide to evaluate the immunogenicity of HPV16 L1/L2 VLPs. Sera of immunized animals were collected two weeks after the third dose.

Western-blot analysis showed that anti-L1/L2 VLP IgG reacted with bands of the proteins L1/L2 purified. These sera detected the presence of one protein similar to protein L1 with approximately 55 kDa and other band with approximately 72 kDa similar to the L2 protein (Fig. 5, column 2, 3). As negative control we used sera from mice immunized with saline that did not react with the same specific proteins (Fig. 5, column 1). Western-blot assays revealed that anti-L1/L2 IgG were able to recognize the HPV 16 L1 and L2 proteins.

**Fig. 5.**
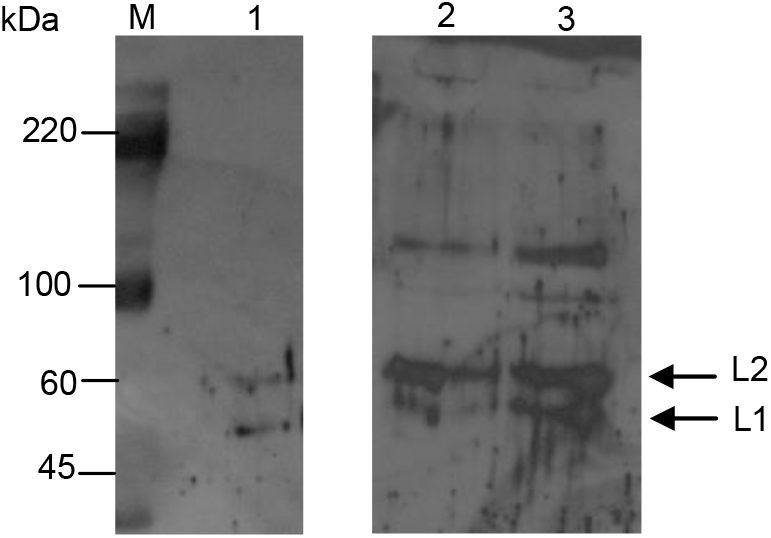
Analysis of the anti-L1/L2 VLP IgG against proteins L1/L2 by Western-blot using membrane incubation with sera samples (pool) from mice immunized with saline (1) and anti-L1/L2 IgG (pool) (2,3), followed by the secondary antibody goat anti-mouse IgG. Molecular mass standards (M)

Antibodies anti-L1 and L2 titers were measured by indirect ELISA assay using pooled sera from animals immunized with protein L1/L2 VLPs or formulation with adjuvant. The sera of immunized animals with formulation L1/L2 VLPs containing adjuvant detected higher titer of anti-L1 IgG (mean= 300), when compared with those induced without adjuvant (mean titer =50). Animals inoculated with saline (control) did not result in detectable levels of either anti-L1 or anti-L2 antibodies. The results between the groups immunized with protein alone (L1/L2 VLPs) and formulations containing the adjuvant were statistically significant (p < 0.05) (Fig. 6A).

**Fig. 6.**
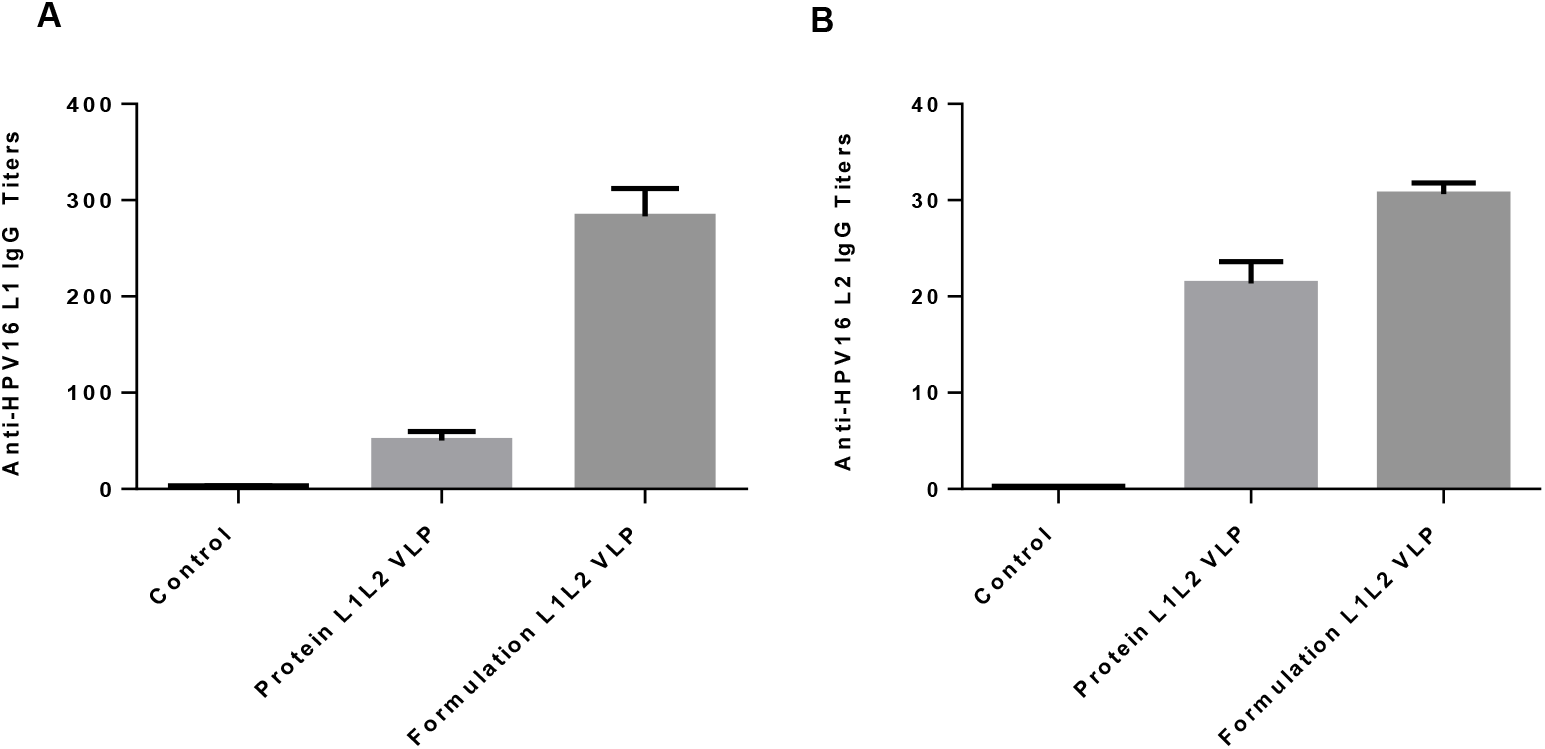
Indirect ELISA analyzes for anti-L1 and anti-L2 IgG after immunization with saline, protein L1/L2 VLP and formulation L1/L2 VLP with adjuvant. Anti-L1 (**A**) and anti-L2 (**B**) IgG titers were detected in pooled sera, two weeks after the last immunization

Antibodies anti-L2 IgG obtained from mice immunized with formulation HPV16 L1/L2 VLP containing aluminum hydroxide showed lower titer (mean=30) as well as the titers of antibodies induced by particles without adjuvant (mean= 20) (Fig. 6B). Statistical analysis showed no significant difference (p > 0.05) between the formulations with or without adjuvant. The control group immunized with saline did not develop antibodies against HPV16 L2 (Fig. 6B).

## Discussion

Human papillomavirus have a great clinical relevance worldwide and has been associated with a wide spectrum of malignancies [17] that justifies the investment in new strategies to control this disease in favor of global public health.

Currently the expression system based on the HEK 293 cell line and derivatives is widely used in research and industrial biotechnology for the production of therapeutic proteins and gene therapy [18]. In this study we used HEK 293-F cells that demonstrated high growth efficiency in serum-free medium with high cell density and viability.

The use of humanized pUF3/L1h (L1) and pUF3/L2h (L2) vectors under the control of the cytomegalovirus promoter (pCMV) has improved the efficiency of HPV16 L1/L2 intracellular protein expression. Several groups have shown that human cell-optimized codons used in different HPV genes increased cell expression [19, 20, 21].

This study demonstrated that the expression system using 293-F cells was well succeeded, presenting high expression efficiency in 48 hours post-transfection, showing that approximately 83% of cells co-transfected with plasmids L1 and L2 expressed the recombinant proteins of interest by quantitative assay.

The overexpression of the HPVs proteins allowed the observation of the intracellular distribution of structural proteins L1 and L2 in cells transfected by indirect immunofluorescence. Both proteins were detected inside the nucleus, dispersed in the cell cytoplasm, which is in accordance with our results described previously [18, 22, 23]. In other study, the expression of HPV11 L1 protein was observed in perinuclear region and in the cytoplasm of mammalian cells, using the immunofluorescence assay [19].

Ultrastructural immunocytochemistry analyses of co-transfected 293-F cells confirmed the expression of the recombinant proteins immune labeling with colloidal gold, identified in the cell nucleus and cytoplasm, which corroborated with results obtained for the immunofluorescence assay, described previously. Furthermore it was also possible to observe the presence of structures of variable sizes, suggesting the formation of virus-like particles (VLPs) composed of two proteins of viral capsid immunolabeled with colloidal gold particles. The self-assembled capsids composed by 72 pentamers of L1 interconnected by L2 were observed by negative staining, corroborating the proposed atomic model for HPV capsids [24].

The purification of the recombinant HPV16 L1/L2 proteins was preceded by the precipitation of the cell lysate with ammonium sulfate, allowing the clarification of the lysate and removal of the contaminants without altering the VLPs conformation [25]. Heparin chromatography was chosen as the method to proceed the purification process. It remains unclear, but heparin has a structure similar to proteoglycan heparan sulfate (HSPG) present on the cell surface, interacting with VLPs in their correct and intact conformation, being an important indicator of the VLPs quality control [25]. The purification conditions are important for maintenance of the VLPs conformation because the recombinant VLPs are unstable and tend to aggregate in solution. To minimize the capsids aggregation and the loss of VLPs during the purification process, we used the stabilizing agent the non-ionic polysorbate surfactant (Tween-80), which can contribute to block the nonspecific bonds of protein [26, 27].

The identity of the purified proteins L1/L2 were demonstrated by Western-blot, which detected the presence of the protein molecular mass approximately 55 kDa, expected for the HPV16 L1 protein [26], recognized by the monoclonal antibody anti-HPV16 (Camvir-1). Other protein was also possible to observe with molecular mass of around 72 kDa, similar with an electrophoretic mobility of the L2 protein in accordance with previous findings [18] and SDS-PAGE analysis showed the molecular mass between 64-78 kDa [28].

The ability of HPV16 L1/L2 VLPs to induce anti-L1 and anti-L2 IgG antibodies in a murine model was demonstrated in Western-blot and ELISA assays.

The specificity of anti-L1/L2 IgG in sera from mice immunized with L1/L2 VLPs against the proteins of viral capsid was confirmed by Western-blot. These sera reacted with the proteins L1 (monomer) and L2 (monomer) in purified proteins obtained from affinity chromatography. Sera of animals used as control, did not recognized the same recombinant proteins.

ELISA assay demonstrated the presence of specific IgG antibodies induced by HPV16 L1/L2 VLPs and the immune responses to both antigens were available. VLP preparation containing adjuvant inoculated in Balb/c mice elicited higher titers of anti-L1 antibodies (mean = 300) than those obtained without adjuvant (mean = 50), suggesting better immunogenicity in formulation containing the adjuvant.

Levels of anti-L2 antibodies induced by the formulations with adjuvant were lower (mean = 30) and similar results of antibody anti-L2 were also found with VLP alone (mean titer = 20). These results indicate that the levels of anti-L2 antibody in mice immunized with HPV16 L1/L2 VLP appear to be lower, as compared with anti-HPV16 L1 IgG. As expected, specific antibodies anti-L1 and anti-L2 were not detected in sera from control animals. These results suggest that the immune response for protein L1 is more robust, when compared to protein L2 in mice immunized with HPV16 L1/L2 VLP formulation.

The immune response of the HPV16 L1 protein in form of L1/L2 VLPs responds dominantly in relation to the specific humoral response of L2, which present a low or undetectable immune response because the L2 protein have being little exposed on the surface of the viral capsid [29, 30]. However, it is known that the amino terminus of the HPV minor capsid protein L2 contains a major cross-neutralization epitope, providing the basis for the development of a broadly protecting HPV vaccine [31, 32].

The results generated in this study suggest that the recombinant HPV16 L1/L2 VLPs may be useful in several studies involving HPV, and also contribute for the development of new vaccine strategies against HPV.

## Ethical approval

Animal studies were performed with the approval of the Ethics Committee of the Butantan Institute (CEUAIB - Protocol Number 923/12).

## Acknowledgments

The authors deep acknowledge Prof. Dr. Martin Müller, German Cancer Research Centre-DKFZ, Heidelberg, Germany for have kindly providing plasmidial vectors; to Prof. Dr. Richard B. Roden, from John Hopkins University, Maryland, USA, for kindly providing the polyclonal antibody anti-L2; to Alexsander Seixas Souza for his technical support in confocal microscope. This work was supported by FAPESP, Butantan Institute and Butantan Foundation.

